# Characterization of pre-analytical blood collection and stabilization parameters to maintain endogenous protein levels for remote blood sampling technology

**DOI:** 10.64898/2026.05.27.728347

**Authors:** Sophie R. Cook, M. Yunos Alizai, Wan-chen Tu, Ingrid Robertson, Xiaofu Wei, Karen Adams, Xiaojing Su, Sanitta Thongpang, Erwin Berthier, Ashleigh B. Theberge

## Abstract

Blood biomarkers are central to monitoring disease progression and evaluating treatment responses, yet traditional venipuncture captures a single physiological snapshot in time and becomes burdensome with repeated sampling. Remote blood self-sampling offers a path toward longitudinal, decentralized monitoring, but maintaining protein integrity from draw to analysis remains a critical challenge. Here, we optimized pre-analytical blood collection and stabilization parameters to maintain protein levels at the time of collection for use with remote sampling technology. First, we optimized blood collection time with Tasso remote self-sampling devices to minimize interference from clotting, finding that a 2.5 min collection time best reduces clot formation while collecting enough blood. Next, we found that Protein Plus, a commercial protein stabilizer, limited hemolysis (a metric for stabilizer efficacy) in venous blood for up to 5 days at 25°C-35°C and for 1 day at 40°C. In addition, we optimized the stabilizer volume and acceptable blood volume range for self-sampling as the stabilizer efficacy is impacted by the stabilizer to blood ratio and collection volume can vary with remote self-sampling devices. Finally, we incubated stabilized blood samples collected via Tasso device at 25°C-35°C for 72 h, mimicking a 2-day shipping period. Using a panel of 21 inflammatory proteins, we found that Protein Plus limited intracellular protein release for various proteins (e.g., VEGF-A, CCL11, and IL-8), inhibited protein degradation for CCL2, and enabled minimal hemolysis. These results support Protein Plus as a viable stabilization strategy for remote blood collection technology targeting longitudinal inflammatory protein monitoring.

---

Blood forms a vast network connecting all organs, serving as a window into the function and interaction of different systems within the body (Figure 1a).^1^ As blood circulates, it carries proteins, RNA, metabolites, and hormones that facilitate inter-organ crosstalk.^2^ Proteins, particularly cytokines and other inflammatory mediators, are of special interest as sensitive indicators of immune status, disease progression, and systemic physiological changes.^3,4^ While single measurements provide a snapshot of biological state, longitudinal sampling (multi-timepoint) captures how these networks evolve over time, offering deeper insight into fundamental human biology.^5^

**Figure 1.**
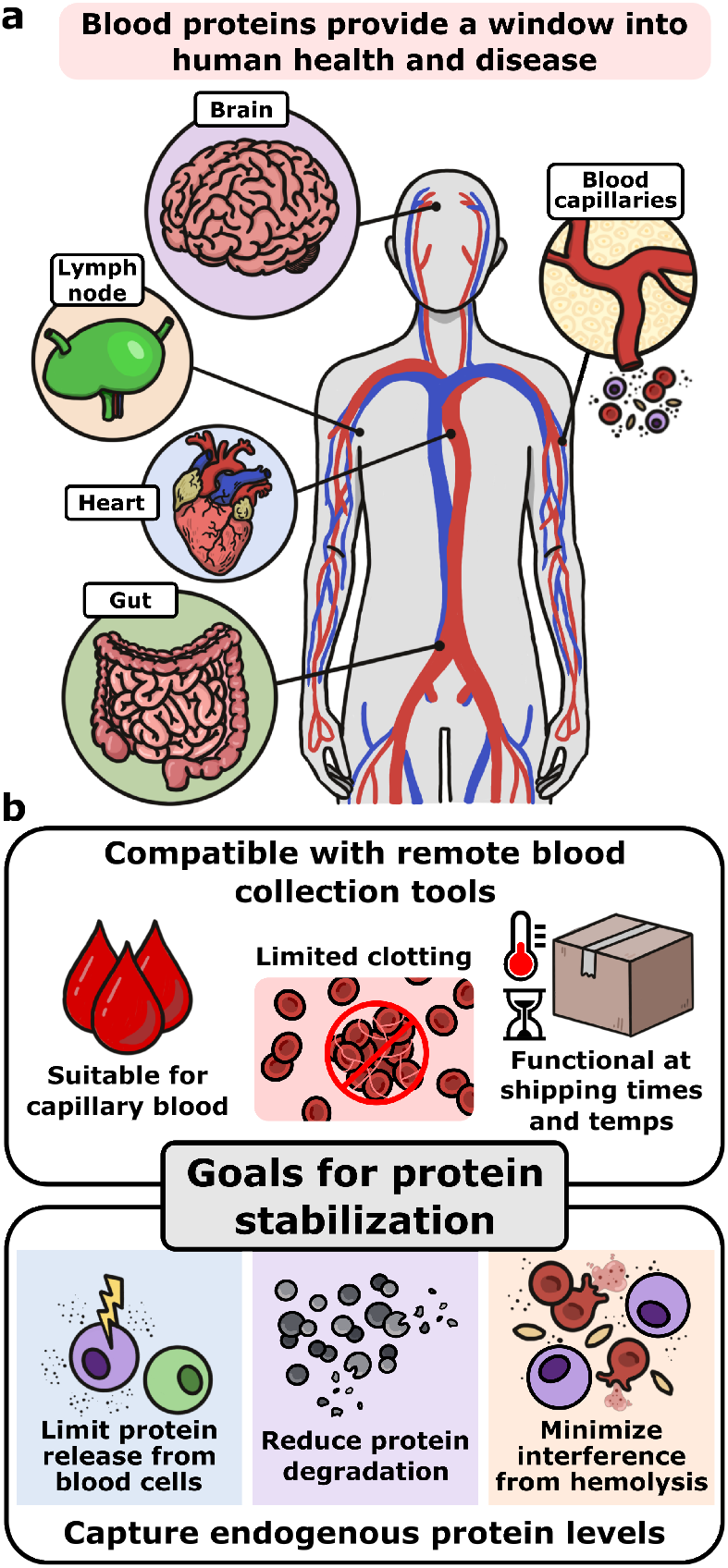
Blood proteins provide a window into human health and disease. (a) Measuring blood proteins can provide a systemic view of various organs in the body and their interactions. (b) The major goals for protein stabilization.

Standard methods for blood testing often rely on venipuncture in clinical settings, typically capturing only a single time point per visit. This introduces significant barriers to longitudinal and large-scale studies where participants must travel repeatedly to clinical facilities, scheduling constraints limit sampling frequency, and logistical demands make recruitment and retention difficult.^6–8^ Collectively, standard testing in the clinic restricts the feasibility of high-frequency sampling and hinders longitudinal studies, making it harder to track transient events, diurnal variation, or long-term trends.

In recent decades, there have been major advancements in remote blood sampling technologies, which have emerged as a promising solution, reducing logistical barriers while enabling high frequency, decentralized collection from participants in their own homes. Several commercially available remote blood sampling approaches have been developed that are not dependent on venipuncture, utilizing either a finger stick or upper arm blood collection stored as dried blood spots (DBS) (e.g., hemaPEN, OneDraw, microfluidic DBS cards)^9–12^ or liquid sample (e.g., Tasso, YourBio Health TAP).^7,13,14^ These devices are widely used for a variety of different biomarkers such as autoantibodies, metabolites, DNA, and inflammatory proteins for applications such as COVID-19, immune-mediated rheumatic disease, organ transplant-related viral infections, wildfire smoke inhalation, and drug pharmacokinetic response.^7–9,15–18^

The aforementioned studies demonstrate a wealth of knowledge on proteins and other analytes from remotely collected blood samples. However, there is an unmet need for systematic characterization of factors affecting protein stability across realistic shipping durations and temperatures. This gap is particularly consequential for cytokine measurement, as cytokines can be released from leukocytes during cell stress or clotting (leading to increased cytokine levels) and many degrade rapidly post-collection without active stabilization (leading to decreased cytokine levels) (Figure 1b).^3^ For venous blood samples collected in the clinic, there are commercially available protein stabilizers such as Protein Plus BCT (now called Nucleic Acid Plus BCT, from Streck) and HEMAcollect Protein BCT (DNA Genotek) that were designed to stabilize proteins and limit hemolysis at ambient temperature for 5-7 days.^19,20^ However, these stabilizers have not been characterized under remote sampling conditions (e.g., self-collected capillary blood, higher temperatures) (Figure 1b). The result is a disconnect between the growing need for decentralized, longitudinal proteomics and the pre-analytical infrastructure required to support it.

Here, we characterized a commercial protein stabilizer, Protein Plus, for use with remote blood sampling technology, addressing a major challenge in remote proteomics. We optimized different parameters such as sample collection time and stabilizer to blood ratio to limit blood clotting and improve stabilizer efficacy in capillary blood at a range of shipping times and temperatures. Finally, we demonstrated successful blood collection and stabilization using remote sampling techniques in a controlled lab environment with 5 participants at shipping times and temperatures. We found that Protein Plus inhibited intracellular protein release, limited protein degradation, and had minimal hemolysis for up to 35°C at 72 h, demonstrating that this commercial stabilizer is compatible with remote sampling methods.

## EXPERIMENTAL

### Participant recruitment

Adult volunteers aged 20-49 years were recruited via word of mouth in Washington State under a protocol approved by the University of Washington Institutional Review Board (STUDY00014133 and STUDY00007868), and written informed consent was obtained from all participants. Each participant was assigned a 3-letter participant code and corresponding participant number that was kept consistent throughout the manuscript (Table S1).

### Whole blood sample collection, stabilization, and processing

Venous blood was collected by venipuncture into 10 mL BD VacutainerTM Plastic Blood Collection Tubes with K2EDTA: HemogardTM Closure (Fisher Scientific). Capillary blood was collected from the upper arm using a Tasso+ or Tasso mini device (Tasso, Inc.) following the manufacture protocol, with the six Tassos as the maximum number of Tasso devices used in one sitting. Briefly, a heat pack (Medline) was applied to the upper arm for 2 min, the area was cleaned with an alcohol pad, and a BD microtainer was attached to the Tasso device before lancet deployment. After collection for 1.5 to 5 min, the tube was sealed and mixed by flicking/inversion. Capillary samples were collected into lavender-top EDTA BD microtainers (Fisher Scientific) for plasma or red-top (no additive) BD microtainers (Fisher Scientific) for serum. All samples in this paper were plasma samples collected into the EDTA tubes except for the clotted controls when optimizing capillary blood collection time. When multiple devices were used per participant, arms were alternated between collections. Blood sampling parameters (time to first blood, collection time, approximate volume) were recorded in Research Electronic Data Capture (REDCap).

Whole blood samples were stabilized using Protein Plus blood collection tube (BCT) (Streck) and was provided in a vacutainer that we removed to use as a reagent. Protein Plus was either added in after sample collection for venous blood or clotted Tasso samples or placed directly in the BD microtainer before sample collection for anticoagulated Tasso samples. Protein Plus BCT has recently been renamed Nucleic Acid Plus BCT but we will refer to it as Protein Plus within this manuscript. After collection and stabilization, plasma and serum were isolated by centrifugation at 1,800 RCF for 15 min. The supernatant was aliquoted into 0.6 ml microcentrifuge tubes, photographed with a smartphone camera against a white background to document plasma or serum color, and stored at –80°C for at least 2 days prior to analysis.

### Protein panel consideration and selection

Proteins were quantified using ProcartaPlex™ multiplex or combinable simplex (single analyte) immunoassay kits (Thermo Fisher Scientific) with a Luminex 200 instrument. We used custom 19plex or 20plex kits that consisted of a range of chemokines, cytokines, growth factors, and other inflammatory proteins (Table 1). These proteins were selected as general biomarkers that may be applicable for a wide range of pathological conditions such as infection, neurodegenerative disease, autoimmunity, cardiovascular disease, and cancer.^4,21–29^ In addition to their relevance for a wide range of inflammatory diseases, we added proteins like interleukin-6 (IL-6), vascular endothelial growth factor-A (VEGF-A), and C-reactive protein (CRP) due to their inclusion on the Vectra test (Labcorp), a blood biomarker panel for rheumatoid arthritis that has clinical interest. Select proteins such as interleukin-8 (IL-8) and interleukin-1 beta (IL-1b) have been found in red blood cell (RBC) lysates and were included as a potential marker for increased hemolysis.^30,31^ We also included four proteins (CD40 ligand (CD40L), IL-1b, IL-8, and VEGF-A) that were previously shown to be stabilized by Protein Plus for up to 5 days at ambient temperature as likely candidates for successful stabilization under shipping times and temperatures.^19^ Finally, we used an established clotting proteins panel (Human Coagulation Panel 3, or Coag. 3) for partially clotted blood samples. More information on the Luminex experimental procedure can be found in the Supplemental Information.

**Table 1.** Inflammatory proteins included in Luminex immunoassay panels.

| Protein | Panel | Protein Type | Relevance |
| --- | --- | --- | --- |
| CCL2 | 19plex, 20plex, 4plex | Chemokine | Chemotaxis in inflammation<sup>4</sup> |
| CCL3 | 19plex | Chemokine | Chemotaxis in inflammation<sup>4</sup> |
| CCL5 | 19plex | Chemokine | Chemotaxis in inflammation<sup>4</sup> |
| CCL11 | 19plex, 20plex | Chemokine | Chemotaxis in inflammation<sup>4</sup> |
| CXCL9 | 19plex, 20plex | Chemokine | Chemotaxis in inflammation<sup>4</sup> |
| IL-1b | 19plex, 20plex | Cytokine | PP* tech. note,<sup>19</sup> RBC lysates,<sup>30</sup> inflammatory disease<sup>21</sup> |
| IL-1RA | 19plex, 20plex | Cytokine | RBC lysates,<sup>30</sup> inflammatory disease<sup>21</sup> |
| IL-4 | 19plex, 20plex | Cytokine | Inflammatory disease<sup>21</sup> |
| IL-6 | 19plex, 20plex, 4plex | Cytokine | Inflammatory disease,<sup>21</sup> Vectra |
| IL-7 | 19plex, 20plex | Cytokine | Inflammatory disease<sup>21</sup> |
| IL-8 | 19plex, 20plex | Cytokine | PP tech. note,<sup>19</sup> RBC lysates,<sup>30</sup> inflammatory disease<sup>21</sup> |
| IL-10 | 19plex, 20plex | Cytokine | Inflammatory disease<sup>21</sup> |
| IL-17A | 19plex, 20plex | Cytokine | RBC lysates,<sup>30</sup> inflammatory disease<sup>21</sup> |
| TNFa | 19plex, 20plex, 4plex | Cytokine | RBC lysates,<sup>30</sup> inflammatory disease<sup>21</sup> |
| IFNg | 19plex, 20plex | Cytokine | RBC lysates,<sup>30</sup> inflammatory disease<sup>21</sup> |
| GM-CSF | 19plex, 20plex | Cytokine | RBC lysates,<sup>30</sup> neurological and inflammatory disease<sup>22</sup> |
| TGFb1 | 19plex, | Cytokine | Inflammatory disease<sup>23</sup> |
| IFNa | 20plex | Cytokine | Inflammatory disease<sup>21</sup> |
| IL-23 | 20plex | Cytokine | Inflammatory disease<sup>21</sup> |
| CD40L | 19plex, 20plex, 4plex | Cytokine | PP tech. note,<sup>19</sup> chronic inflammation and autoimmunity<sup>24</sup> |
| VEGF-A | 19plex, 20plex | Cytokine, Growth Factor | PP tech. note,<sup>19</sup> RBC lysates,<sup>30</sup> inflammation,<sup>25</sup> Vectra |
| Ferritin | 20plex | Cytokine, Iron storage | Inflammation, autoimmunity, infectious disease<sup>27</sup> |
| Calcitonin | 20plex | Hormone | Cancer and autoimmunity<sup>26</sup> |
| CRP | Simplex | Inflammatory marker | Clinical blood biomarker,<sup>28</sup> Vectra |
| vWF | Coag. 3 | Coagulation factor | ↑ with clotting<sup>32</sup> |
| Factor IX | Coag. 3 | Coagulation factor | ↓ with clotting<sup>32</sup> |
| Protein C | Coag. 3 | Anticoagulation factor | ↓ with clotting<sup>32</sup> |
| Protein S | Coag. 3 | Anticoagulation factor | ↓ with clotting<sup>32</sup> |

\*PP = Protein Plus

### Optimization of Tasso blood collection time to reduce blood clotting and minimize changes in protein concentration

7 healthy participants (6 female and 1 male, P1-7) were recruited for this study. Each participant collected capillary blood on two occasions three weeks apart, using Tasso+ devices first and Tasso mini devices three weeks later. For each Tasso collection, an EDTA BD microtainer was pre-loaded with 50 µL of Protein Plus stabilizer. Collection times per tube were 1.5 min, 2.5 min, and 5 min (or until the blood volume reaches 1 mL, with 3 min being the lowest observed time) (Table S2). After collection, the collection tubes were sealed and mixed by flicking/inversion. For 3 participants (P1-3), an additional sample was collected for 3-5 min in a red top BD microtainer (no additive) and allowed to clot for 30 min at room temperature before adding 50 µL of Protein Plus and inverting to mix. Plasma and serum were then isolated via centrifuge as described above and stored at -80°C for at least 2 days prior to quantification via Luminex.

To determine the optimal blood collection time, estimate blood volumes were measured via pipette and any clots observed when pipetting were noted. Proteins were quantified using a custom 19-plex kit and a 4-plex Human Coagulation Panel 3 kit (Table 1). Samples were diluted 1:4 in platinum buffer prior to plate preparation based on prior results (Figure S1). When incubating the sample with the beads, we used a 2 h incubation period with 7 standards. The raw data were analyzed using the ProcartaPlex Analysis App (Thermo Fisher Scientific) using a five-parameter logistic standard curve, and the concentration and/or fold change were plotted in Prism 10 (GraphPad). Conditions where the bead count was below 30 beads were excluded. When the reported concentration was listed as either <OOR (outside of range) or the extrapolated value was below the lower limit of quantitation (LLOQ), the value was replaced with the LLOQ.

### Impact of protein stabilizer on hemolysis at a range times and temperatures using venous blood

2 healthy participants (2 females, P1 and P3) were recruited for this study. Venous blood was collected from each participant by venipuncture into EDTA-coated vacutainer tubes as described above. Whole blood from each participant was pooled and aliquoted into 100 µL volumes in 0.6 mL microcentrifuge tubes, followed by addition of 5 µL Protein Plus stabilizer. Tubes were sealed and mixed by flicking/inversion. Aliquots were then incubated at 25°C, 35°C, 40°C, and 45°C for 24 h, 48 h, 72 h, and 120 h with a 0 h control. Following incubation, plasma was isolated as described above before the samples were photographed and stored at –80°C.

### Varied protein stabilizer ratios with venous blood at shipping times and temperatures

1 healthy participant (male, P8) was recruited for this study. Venous blood was collected by venipuncture into an EDTA-coated vacutainer tube as described above. Protein Plus stabilizer was added to whole blood at varying stabilizer to blood ratios: 0 (no stabilizer), 1:3, 1:5, 1:10, 1:15, 1:20, and 1:30 (Table S3). The 1:3 ratio reflects the minimum expected collection volume on a Tasso device (150 µL blood) with 50 µL stabilizer pre-loaded per microtainer; the 1:20 ratio was selected based on the manufacturer’s recommendation (Streck). Blood stabilizer mixtures were aliquoted in duplicate into 0.6 mL microcentrifuge tubes and incubated at 25°C, 35°C, and 40°C for 0 h, 48 h, and 72 h. Following incubation, plasma was collected and imaged as described above, then stored at –80°C.

### Protein stabilizer efficacy at shipping times and temperatures using capillary blood

5 healthy participants (2 female and 3 male, P2, P8-11) were recruited for this study. Each participant collected capillary blood using six Tasso mini devices with EDTA BD microtainers, three pre-loaded with 50 µL of Protein Plus and three without. Blood was collected either until 2.5 min had passed or until 500 µL of blood was collected, whichever came first (Table S4). After collection, tubes were sealed and mixed by flicking/inversion. Samples from matching conditions were pooled and split into 100-150 µL aliquots in 0.6 mL centrifuge tubes. Aliquots were then incubated in duplicate at 25°C and 35°C for 72 h to simulate shipping conditions with 0 h controls. Following incubation, plasma was collected and photographed as described above before storage at –80 C for at least 2 days before analysis. Proteins were quantified using a custom 20-plex kit and a CRP simplex kit (Table 1). For the 20-plex kit, the plate was prepared with 8 standards and samples were diluted 1:4 in platinum buffer prior to plate preparation. The CRP simplex kit was prepared separately with 7 standards and samples were diluted 1:500 in UAB. For the first incubation step with plasma samples and beads, the 20-plex kit was incubated overnight at 5°C and the CRP kit was incubated for 2 h at ambient temperature. The raw data were analyzed as described above. The fold change was calculated for each condition by first averaging the duplicate concentrations and then dividing the average by the 0 h control concentration.

## RESULTS AND DISCUSSION

### Considerations for protein stabilization in remote sampling

To start, we selected the commercially available Tasso blood collection device as our model remote sampling tool, a device that utilizes a lancet with added vacuum to collect capillary blood from the upper arm. We tested two models of the Tasso device with varying lancet sizes: the Tasso+ device with a lancet length of 3.8 mm and depth of 0.8 mm and the Tasso mini with a lancet length of 1.8 mm and depth of 1.6 mm. While we are using these Tasso devices for this study, our findings are translatable to a range of different remote sampling technologies with similar capillary blood collection mechanisms.

For the stabilizer characterization described here, we use the commercial stabilizer, Protein Plus, which consists of an anticoagulant and a cell preservative to prevent interference from blood cell proteins that would otherwise be released during hemolysis, platelet activation, or immune cell stress. Here, we added Protein Plus directly to the collection tube before sampling, resulting in little change for the original Tasso blood collection workflow. This method was straightforward for collecting samples in a controlled lab environment. We are currently exploring methods to easily introduce the stabilizer to the blood sample similar to technology we developed for RNA stabilization to facilitate user-friendly at-home use.^14,33,34^

There is a wide range of available methods to quantify blood proteins such as mass spectrometry,^35^ proximity extension assays (PEA) (e.g., Olink),^36^ and multiplex immunoassays like Luminex^37^ or the single-molecule assay (Simoa).^38^ We selected the Luminex multiplex immunoassay as our method to quantify our selected proteins of interest since it was relatively low-cost to measure 10-20+ specific proteins within our own lab space. With the Luminex assay, we chose to focus on a subset of 19 to 21 inflammatory proteins that are prevalent in autoimmune disease, long COVID, infection, cancer, neurodegenerative disease, cardiovascular disease, and more.^21–29^ We hypothesize that the commercial stabilizer Protein Plus would be compatible with other quantification methods and has been characterized with both Luminex assays and Olink assays by the vendor.^19^

### Reduced Tasso collection time limits blood clotting and minimizes impact on downstream protein quantification

When testing the pre-analytical blood collection workflow, we needed to consider both the collection conditions (e.g., collection time, lancet size) and the sample matrix (i.e., plasma or serum). To measure blood proteins, the liquid fraction is separated from the cellular fraction of the blood. If the separation occurs without clotting, the liquid fraction is referred to as plasma, whereas the liquid portion after clotting is referred to as serum.^39^ While multiplexed immunoassays like Luminex are compatible with plasma and serum samples, these complex matrices may inhibit the detection of proteins, especially cytokines that are typically present in low concentrations (<10 pg/mL). We used a 4plex kit of combined single analyte kits for key proteins such as IL-6, tumor necrosis factor-alpha (TNFa), and C-C motif chemokine ligand 2 (CCL2), to optimize spiked-in protein recovery by diluting the complex plasma matrix in platinum buffer at varying ratios, finding that a 1:4 dilution was sufficient to reduce matrix effects (Figure S1).

Protein Plus was designed for and exclusively tested on plasma (i.e., no clotting).^19^ While this stabilizer may be effective for proteins in serum, it would be challenging for the stabilizer to penetrate a clot in coagulated blood to prevent effects such as immune cell stress that may impact the endogenous protein levels in the sample.^32^ However, it is challenging to obtain a standard plasma sample from at-home blood collection devices like Tasso due to the inherent nature of capillary blood collection as the blood may be partially clotted by the time it reaches the anticoagulant-coated collection tube. To determine the ideal blood collection time and the potential impact of blood clotting on endogenous protein levels, we collected samples at varied collection times (3-5 min (the standard collection time), 2.5 min, and 1.5 min) with devices of different lancet sizes (Tasso+ and Tasso mini) for 7 participants, with an additional clotted control for 3 participants (Figure 2a). We measured protein concentration using a custom 19plex Luminex immunoassay consisting of inflammatory cytokines, chemokines, growth factors, and other proteins of interest in addition to a 4plex panel of coagulation factors.

**Figure 2.**
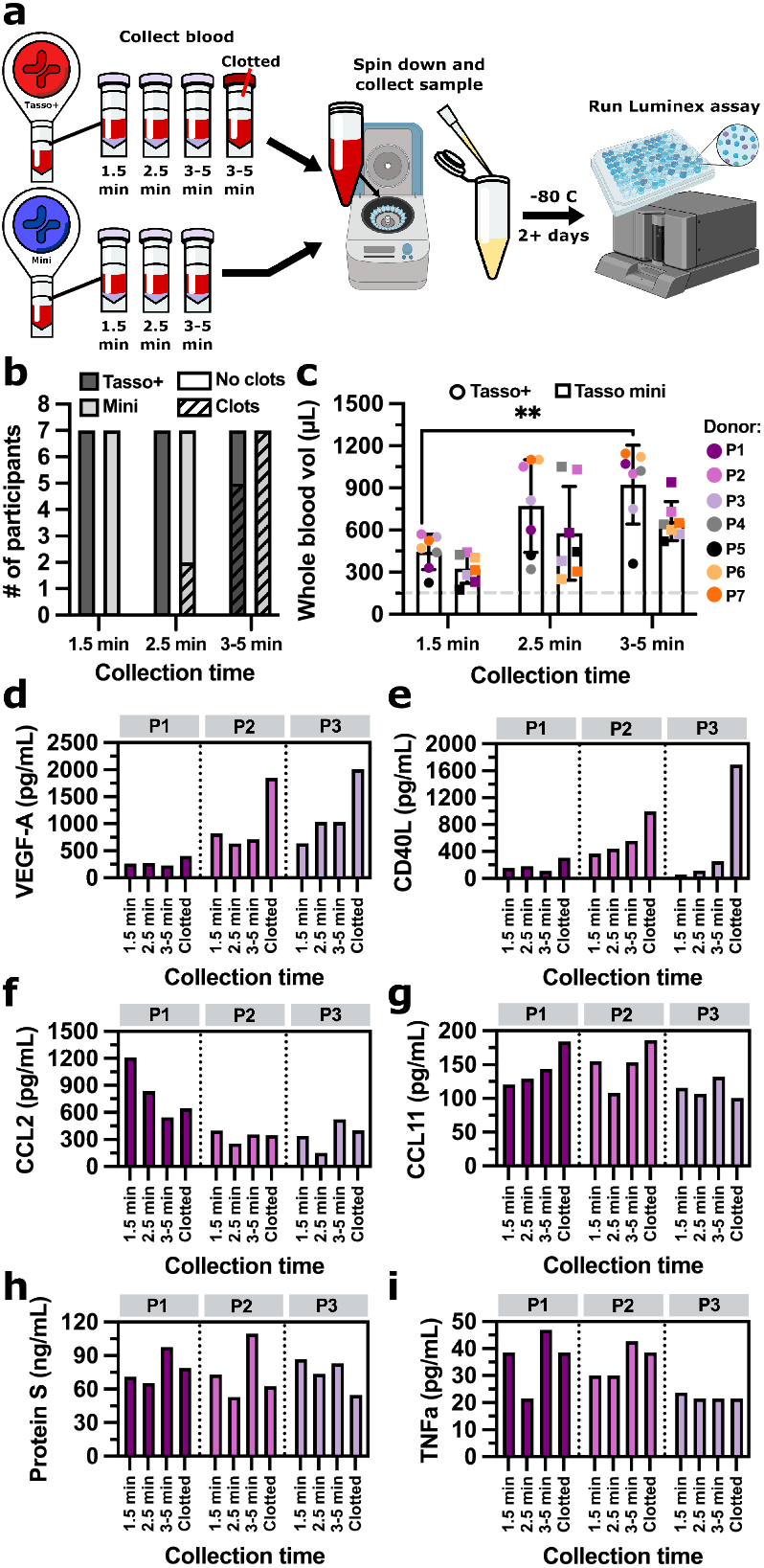
Impact of blood collection time on blood clotting and endogenous protein levels. (a) Samples were collected at different times via the Tasso+ and Tasso mini collection device. (b) The number of samples for both the Tasso+ and mini where clots were observed at different collection times. (c) Estimated whole blood volume for all 7 donors (P1-P7) for Tasso+ and Tasso mini devices. The minimum volume preferred for the downstream assay (150 µL) is noted with a grey dashed line. Each point represents a single sample collected with a Tasso device. Results were compared using a two-way ANOVA with Tukey post-hoc tests (n = 7). All comparisons are not significant (p > 0.05) unless noted otherwise. ** indicates p < 0.01. (d-h) The concentrations of VEGF-A, CD40L, CCL2, CCL11, Protein S, and TNFa for the Tasso+ for 3 donors (P1, P2, and P3). Each bar represents plasma or serum from a single Tasso collection.

When testing different blood collection times, we found that a reduced collection time of 1.5 min and 2.5 min had fewer observable clots (Figure 2b) and still produced above 150 µL of blood, which is sufficient for downstream analysis (Figure 2c). We found that some sensitive cytokines such as IL-6 were below the assay limit of detection (data not shown), while 9 proteins such as C-C motif chemokine ligand 11 (CCL11), VEGF-A, TNFa, Protein S, and CCL2 were within the assay range (Figure 2d-i). When looking at results that include a clotted control from the first three participants, there were notable trends for three proteins of interest. For VEGF-A and CD40L, the protein concentration increased when the sample was clotted, a trend especially evident with two of the three participants (Figure 2d, e). For CCL2, one participant (P1) saw a decrease in protein concentration as collection time increased (Figure 2f). We also saw an increase in CCL11 for the same participant (P1) with variable results for the other two participants (Figure 2g). For Protein S, the anticoagulation factor involved in the clotting cascade, the 1.5 min and 2.5 min collection times were relatively consistent with increased variability at 3-5 min (Figure 2h). The protein concentration across blood collection times for TNFa was variable, and with limited replicates, no conclusions could be drawn for this protein (Figure 2i). *Based on these results, we selected 2*.*5 min as our goal collection time for the Tasso+ and mini to both minimize clotting to maintain endogenous protein levels while also collecting enough blood volume consistently*. While this experiment was conducted using a Tasso blood collection device, these results can be generalizable for other remote sampling devices where sample collection parameters like collection time can be optimized to limit blood clotting.

### Hemolysis of venous blood at a range of shipping times and temperatures

As remote samples undergo shipping to return the samples to the lab for downstream testing, it is critical that the commercial stabilizer remains effective at times and temperatures that samples may experience during shipment. While Protein Plus has been marketed as effective to inhibit hemolysis and stabilize select plasma proteins at room temperature (∼25°C) for up to 5 days, shipping temperatures can be much higher, especially in summer months, increasing up to 40°C-45°C. Here, we tested the effectiveness of Protein Plus to inhibit hemolysis at a range of times (24-120 h) and temperatures (25°C-45°C) with a 0 h control across two participants. We compiled images of plasma at each condition in triplicate (Figure 3).

**Figure 3.**
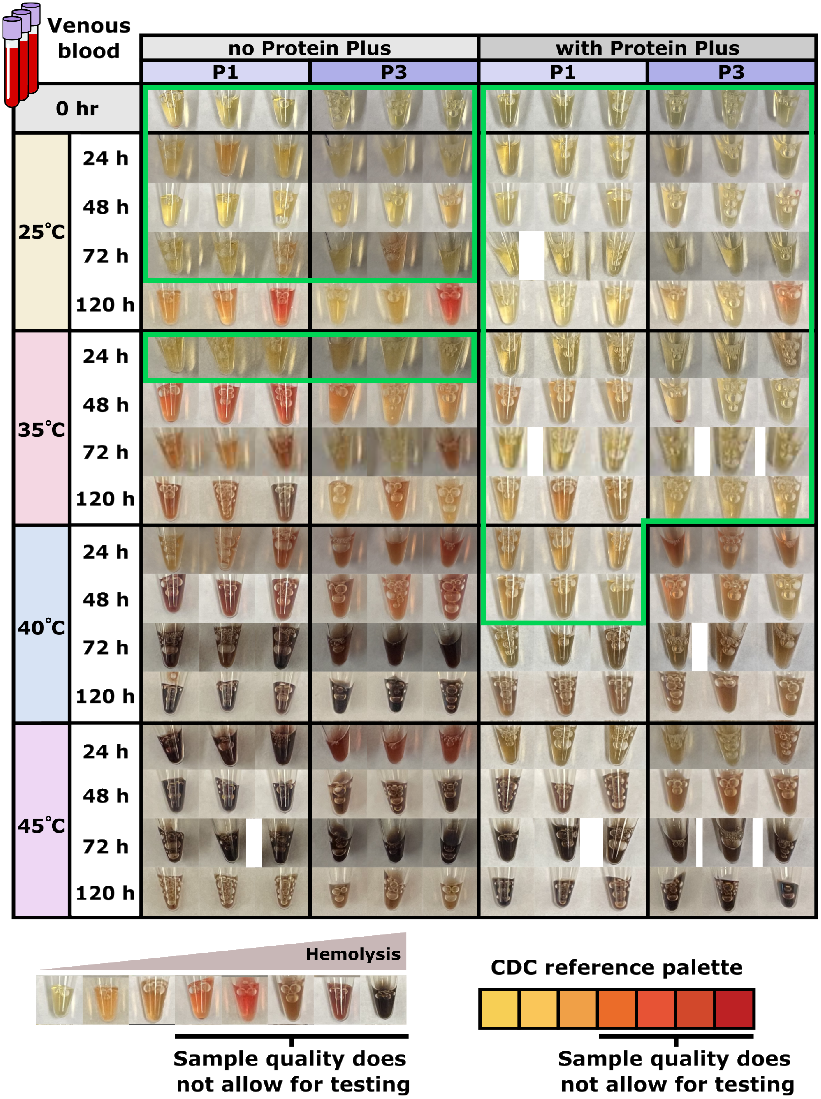
Protein stabilizer inhibits hemolysis in venous blood. Images of plasma arrayed in a heat map for each condition for 2 participants (P1 and P3). Venous blood was stabilized at the recommended stabilizer to blood ratio of 1:20. The conditions where the plasma looks to be within the acceptable hemolysis range are boxed in green. The CDC reference palette and sample quality restrictions were taken from reference 19.

We used plasma color here as a qualitative readout for hemolysis,^40^ where clear/yellow plasma color indicates little to no hemolysis and orange/pink to red plasma color indicates increasing hemolysis levels. At higher temperatures, the hemoglobin can oxidize, resulting in light brown to dark brown plasma color, where we would expect significant interference in downstream analysis. We are using the CDC plasma color palette as a reference for sample quality restrictions, though this palette was designed for use with general lab testing and not our specific protein quantification method; we measure these proteins directly in experiments later in the manuscript.

With Protein Plus present, the samples were usable at 25°C and 35°C for 120 h for both participants and 40°C for 48 h for one participant (Figure 3). Without a stabilizer, the samples had limited hemolysis only at 25°C for 72 h and 35°C for 24 h. The samples at 45°C, even at 24 h, were not usable with or without the stabilizer. *These results indicate that Protein Plus effectively inhibits hemolysis at shipping times and temperatures but would not be effective in high temperature climates without more temperature control*.

### Protein stabilizer hemolysis inhibition is impacted by stabilizer to blood ratio

Commercial stabilizers like Protein Plus have recommended ratios to use with venous draws where the final blood collection volume is known. For remote sampling, it is challenging to know what volume of stabilizer to use because the volume of blood collected is variable, resulting in a wide range of possible stabilizer to blood ratios (Figure 4a). In preliminary work, we saw increased hemolysis levels with capillary blood collected using a Tasso device even at 25°C for 72 h (Figure 4b, methods in Supplement), a condition where we saw little to no hemolysis with venous blood where the ratio was controlled (Figure 3). To measure the impact of protein stabilizer to blood ratio on hemolysis, we added a range of volumes of Protein Plus to a constant volume of venous blood. We selected the different stabilizer volumes based on the range of possible ratios when collecting blood with the Tasso device with a constant 50 µL of stabilizer and blood volumes of 150 µL to 1 mL, ranging from 1:3 to 1:30 of stabilizer to blood with 1:20 being the recommended ratio from the vendor.

**Figure 4.**
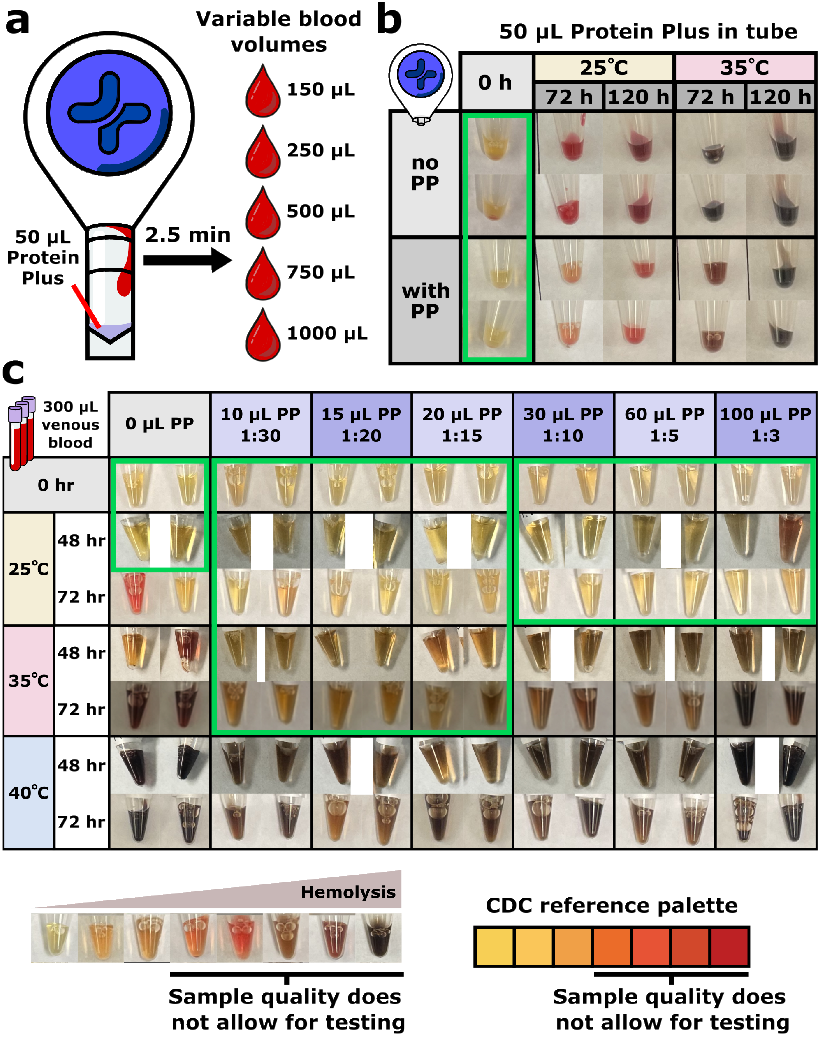
The ratio of stabilizer to blood impacts hemolysis inhibition. (a) Since the final blood volume collected via Tasso varies from 150-1000 µL, the resulting ratio of Protein Plus (PP) stabilizer to blood varies as the stabilizer volume remains constant. (b) Images of plasma from capillary blood arrayed in a heat map for each condition for 1 participant (P5). Each tube was preloaded with 50 µL of Protein Plus before collection. The conditions where the plasma looks to be within the acceptable hemolysis range are boxed in green. (c) Images of plasma from venous blood with various stabilizer (PP) ratios arrayed in a heat map for each condition for 1 participant (P8). The conditions where the plasma looks to be within the acceptable hemolysis range are boxed in green. The CDC reference palette and sample quality restrictions were taken from reference 19.

After incubating the samples at 25°C, 35°C, and 40°C for 48 h and 72 h, we found that the stabilizer ratio does impact the hemolysis levels. This was especially evident at 35°C, where the 1:15, 1:20, and 1:30 ratios showed less hemolysis and was within the acceptable plasma color range for CDC guidelines compared to 1:10, 1:5, and 1:3 (Figure 4c). *Based on these results, we decided to decrease the volume of stabilizer that we add to the blood collection tube to 20 µL and have a new acceptable blood volume range (250-500 µL)*. As a result, our potential ratios now range from 1:12.5 (250 µL of blood) to 1:25 (500 µL of blood).

Protein stabilizer inhibits intracellular protein release and limits protein degradation for select proteins at relevant shipping times and temperatures. Finally, we sought to model the shipping conditions of remote sample collection by collecting capillary blood with Tasso blood collection devices followed by a 72 h incubation at 25°C and 35°C, mimicking the length and typical temperatures for an over-night shipping period.^33^ We collected whole blood from five healthy participants using multiple Tasso mini devices that were then pooled before aliquoting the samples for incubation, plasma isolation, and downstream protein quantification (Figure S2). *Using the optimized stabilizer volume and blood volume range, we found that all five participants had limited hemolysis at both 25°C and 35°C after the 72 h incubation with stabilizer present, while we observed hemolysis present at 35°C with no stabilizer (Figure 5a)*.

**Figure 5.**
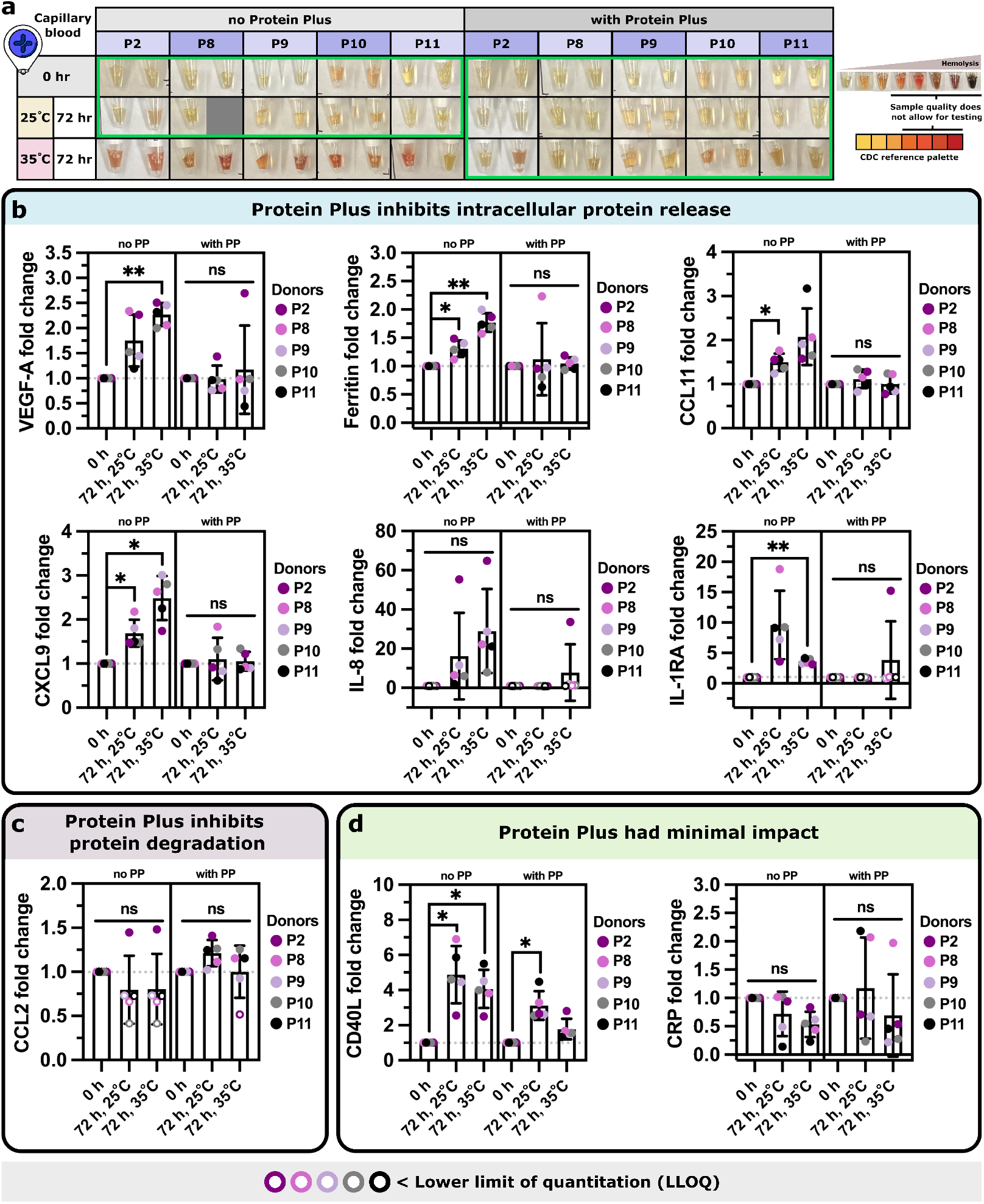
Protein Plus limits hemolysis and maintains endogenous protein levels for select proteins under relevant shipping times and temperatures. (a) Images of plasma from capillary blood with or without Protein Plus (PP) stabilizer arrayed in a heat map for each condition for 5 participants (P2, P8, P9, P10, and P11). Missing replicates are represented as a grey box. The conditions where the plasma looks to be within the acceptable hemolysis range are boxed in green. The CDC reference palette and sample quality restrictions were taken from reference 19. (b) The addition of the protein stabilizer inhibits intracellular protein release for VEGF-A, ferritin, CCL11, CXCL9, IL-8, and IL-1RA. (c) The addition of the protein stabilizer inhibits protein degradation for CCL2. (d) The addition of the protein stabilizer had minimal impact on CD40L and CRP. For (b-d), each point represents the average fold change of two replicates compared to 0 h control for 5 participants (P2, P8, P9, P10, and P11). A fold change of 1 (no change) is marked on all graphs with a grey dotted line. Replicates that were below the lower limit of quantitation (LLOQ) were replaced with the LLOQ value per Luminex recommendation before averaging the replicates and calculating the fold change. Points where either one or both replicates were below the LLOQ are noted as hollow points. Results were compared using a one-way ANOVA with repeated measures and Tukey post-hoc tests (n = 5). All comparisons are not significant (ns, p > 0.05) unless noted otherwise. ** indicates p < 0.01, * indicates p < 0.05, ns indicates p > 0.05.

Next, we quantified a range of inflammatory proteins using a custom 20-plex Luminex kit paired with a single-analyte CRP kit (Table 1). The new 20plex kit includes many of the same proteins of interest as the previous 19plex kit such as VEGF-A and IL-8 while some proteins with inconsistent results (e.g., C-C motif chemokine ligand 3 (CCL3)) were replaced with other proteins of interest such as ferritin and calcitonin. To standardize the changes in protein levels across all five participants, we reported the fold change of each protein compared to the 0 h control within each participant rather than the raw concentration. Of the 21 proteins measured, 11 proteins were below the assay range for most if not all samples, including many sensitive cytokines like IL-6 and TNFa. As this experiment was done with five healthy participants, this was to be expected with some proteins on the panel as we predict that many of these proteins would be significantly higher in instances of inflammation.

Of the remaining 9 proteins with quantifiable results, we found that for VEGF-A, ferritin, CCL11, chemokine ligand 9 (CXCL9), IL-8, and interleukin-1 receptor antagonist (IL-1RA), the protein stabilizer inhibited intracellular protein release, with protein levels staying relatively constant with stabilizer present while the samples with no stabilizer increased with the incubation (Figure 5b). We saw an especially large increase in intracellular protein release without the stabilizer present for IL-8, a cytokine often found in reservoirs in RBCs,^30,31^ with a 60–70-fold change with increased incubation time and temperature, indicating potential RBC lysis that was not visible in the plasma color alone. For CCL2, we saw that the stabilizer inhibited protein degradation with a relatively constant fold change for four of the participants with stabilizer present, with decreasing protein levels with no stabilizer present (Figure 5c). For CD40L and CRP, the stabilizer had minimal impact on the protein levels with increased time and temperature (Figure 5d). *In summary, we demonstrated that the commercial stabilizer, Protein Plus, was beneficial to minimize hemolysis and maintain endogenous protein levels for select inflammatory proteins (VEGF-A, ferritin, CCL11, CXCL9, IL-8, IL-1RA, and CCL2) at shipping temperatures and times (25°C and 35°C for 72 h)*. We expect that this stabilizer may not be effective for all proteins, as we saw with CD40L and CRP, so we recommend careful characterization when using this stabilizer for new protein panels.

## CONCLUSION

Here, we demonstrated that Protein Plus effectively stabilizes a panel of inflammatory proteins in whole blood by suppressing intracellular protein release, inhibiting extracellular protein degradation, and limiting hemolysis for up to 72 hours at temperatures up to 35°C. While current remote shipping conditions in the US have been shown to produce transient temperature spikes reaching into the low to mid 40s°C (range 35.4-45.1°C across 14 US states in summer shipping),^34^ and up to 41°C during brief courier transit in hot climates such as Doha, Qatar,^33^ these excursions are typically short lived. Given that Protein Plus maintained protein stability at 35°C across the full 72-hour duration tested here, these short-lived excursions may not compromise sample integrity under real world shipping conditions. In regions that regularly experience more extreme ambient temperatures, such as tropical climates or the Middle East, the incorporation of phase change materials into sample packaging could serve as an additional mitigation strategy to buffer against extreme temperature fluctuations during transit.

Importantly, we demonstrated compatibility between Protein Plus and Tasso self-sampling devices, establishing a foundation for at-home longitudinal blood protein collection mitigating the risk of protein degradation or artifactual signal from cellular release. These findings represent a meaningful step toward decentralized biomarker monitoring outside traditional clinical settings. However, Protein Plus did not fully suppress hemolysis under all tested conditions, which may introduce confounds for hemolysis-sensitive analytes and warrants further investigation. Additionally, the current format (20 µL of Protein Plus directly in the collection tube) may not represent the optimal configuration for reliable at home collection. Therefore, we are currently exploring user-friendly alternative methods of introducing the stabilizer to the blood sample, drawing on approaches established in our homeRNA platform. We also aim to validate this approach under real world conditions with participants collecting samples remotely with return shipping to the lab. Longer term integration with our homeRNA platform presents an opportunity to develop a multiomics platform capable of simultaneously stabilizing proteins, RNA, and additional analytes of interest from a single, convenient self-collected blood sample.

## Supporting information

Supplemental information including raw data description, supplemental methods, supplemental figures, and supplemental tables.

## ASSOCIATED CONTENT

### Supporting Information

The Supporting Information is available free of charge at [insert link].

Description of raw data available, additional methods and figure on sample matrix effects experiment, table on participant numbers and codes, summary tables of Tasso blood collection data (PDF).

Raw data files for all Luminex immunoassay experiments (CSV, XLS, PDF).

## AUTHOR INFORMATION

### Present Addresses

Sophie R. Cook – Department of Chemistry, University of Washington, Seattle, Washington 98195; United States;

M. Yunos Alizai – Department of Chemistry, University of Washington, Seattle, Washington 98195; United States;

Wan-chen Tu – Department of Chemistry, University of Washington, Seattle, Washington 98195; United States;

Ingrid Robertson – Department of Chemistry, University of Washington, Seattle, Washington 98195; United States;

Xiaofu Wei – Department of Chemistry, University of Washington, Seattle, Washington 98195; United States;

Karen Adams – Department of Chemistry and Institute of Translational Health Sciences, School of Medicine, University of Washington, Seattle, Washington 98195; United States;

Xiaojing Su – Department of Chemistry, University of Washington, Seattle, Washington 98195; United States;

Sanitta Thongpang – Department of Chemistry, University of Washington, Seattle, Washington 98195; United States;

### Author Contributions

All authors have given approval to the final version of the manuscript.

### Notes

The authors declare the following potential competing financial interest(s): SRC, MYA, ST, EB, and ABT filed patent 64/054,742 through the University of Washington on multiomics remote blood sampling tools. EB and ABT filed patent 17/361,322 (Publication Number: US20210402406A1) and patent 63/571,012 through the University of Washington on homeRNA and a related technology. ABT, ST, XS reports filing multiple patents through the University of Washington. ABT received a gift to support research outside the submitted work from Ionis Pharmaceuticals. SRC reports filing multiple patents through the University of Virginia on organs-on-chip devices and related technology. EB and ST have ownership in Salus Discovery, LLC, and Tasso, Inc., and EB is employed by Tasso, Inc. Tasso, Inc. developed the blood collection systems used in this publication. Technologies from Salus Discovery, LLC are not included in this publication. EB, ST, and ABT have ownership in Seabright, LLC, which will advance new tools for diagnostics and clinical research including homeRNA and other stabilizing solutions, and EB and ST are partially employed by Seabright, LLC. The terms of this arrangement have been reviewed and approved by the University of Washington in accordance with its policies governing outside work and financial conflicts of interest in research.

## ACKNOWLEDGMENTS

This publication was supported by Schmidt Sciences, LLC and the National Institutes of General Medical Sciences (NIGMS) of the National Institute of Health (NIH) under award number R35GM128648. The REDCap used for human subjects consent and enrollment was supported by the Institute of Translational Health Sciences (ITHS) under the National Center for Advancing Translational Science of the NIH under award number UL1TR002319. The content is solely the responsibility of the authors and does not necessarily represent the views of the NIH or other funding bodies.

We would like to thank Damon Cook and David Soltys from Thermo Fisher for training, consultations, and discussions on the Luminex immunoassay protocol and analysis. We would like to thank Nidhi Ghadge for assistance collecting capillary blood samples and recording sampling information. We would like to thank Meredith Harris for editorial support. Finally, we would like to thank the study participants without whom this work would not be possible.

## Table of Contents Figure

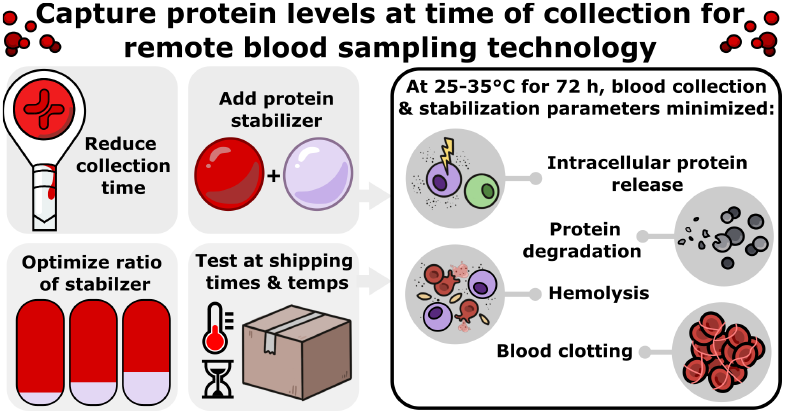

