## Supplemental information including raw data description, supplemental methods, supplemental figures, and supplemental tables. for "Characterization of pre-analytical blood collection and stabilization parameters to maintain endogenous protein levels for remote blood sampling technology"

Electronic Supplementary Information for

**Contents:**

- Description of Luminex raw data
- Supplemental Methods
- Supplemental Figures
- Supplemental Tables

### I. DESCRIPTION OF LUMINEX RAW DATA

We included the raw data from all three Luminex experiments done within the manuscript, all attached separately. This includes the standard curves, upper and lower limit of quantitation (ULOQ and LLOQ), net mean fluorescent intensity (net MFI), and extrapolated concentration for each protein within each assay. We also included the CSV files directly from the Luminex 200 instrument for each assay so the data can be re-analyzed with the ProcartaPlex analysis app or analysis method of choice using the kit catalogue and lot numbers provided in Table S5.

**File names and contents:**

- Figure 2 file names and descriptions
    - Fig 2_clotting_19plex_raw data.csv
        - Raw data file generated by Luminex 200 instrument
    - Fig 2_clotting_19plex_standard curves.pdf
        - PDF following analysis via Procartaplex Analysis App showing standard curves
    - Fig 2_clotting_19plex_organized data.xlsx
        - Spreadsheet of extrapolated concentration and net MFI. The concentration results have been labeled and organized. A key is included to define labels.
    - Fig 2_clotting_Coag 3_raw data.csv
        - Raw data file generated by Luminex 200 instrument
    - Fig 2_clotting_Coag 3_standard curves.pdf

        - PDF following analysis via Procartaplex Analysis App showing standard curves
    - Fig 2_clotting_Coag 3_organized data.xlsx
        - Spreadsheet of extrapolated concentration and net MFI. The concentration results have been labeled and organized. A key is included to define labels.
- Figure 5 file names and descriptions
    - Fig 5_Tasso temp stress_20plex_raw data.csv
        - Raw data file generated by Luminex 200 instrument
    - Fig 5_Tasso temp stress_20plex_standard curves.pdf
        - PDF following analysis via Procartaplex Analysis App showing standard curves
    - Fig 5_Tasso temp stress_CRP_raw data.csv
        - Raw data file generated by Luminex 200 instrument
    - Fig 5_Tasso temp stress_CRP_standard curves.pdf
        - PDF following analysis via Procartaplex Analysis App showing standard curves
    - Fig 5_Tasso temp stress_20plex and CRP_organized data.xlsx
        - Spreadsheet of extrapolated concentration and net MFI. The concentration results have been labeled and organized. A key is included to define labels.
- Figure S1 file names and descriptions
    - Fig S1_matrix effects_raw data.csv
        - Raw data file generated by Luminex 200 instrument
    - Fig S1_matrix effects_CD40L_standard curves.pdf
        - PDF following analysis via Procartaplex Analysis App showing standard curves
    - Fig S1_matrix effects_IL6 TNFa CCL2_standard curves.pdf
        - PDF following analysis via Procartaplex Analysis App showing standard curves
    - Fig S1_matrix effects_organized data.xlsx
        - Spreadsheet of extrapolated concentration and net MFI. The concentration results have been labeled and organized. A key is included to define labels.

### II. SUPPLEMENTAL METHODS

**Protein quantification using Luminex immunoassays.** All plasma and serum samples were thawed on ice before use. Assay reagents were brought to room temperature and prepared according to the manufacturer's instructions. For all kits (excluding CRP simplex kit), the universal assay buffer (UAB) was replaced with platinum buffer (ProcartaPlex$^{TM}$ Platinum Assay Buffer for Plasma Samples, Thermo Fisher Scientific), including standard preparation and serial dilution. Platinum buffer has been recommended by the vendor to use as a diluent to reduce matrix effects from plasma and serum samples. Since the CRP simplex kit required a significant dilution of 1:500, we used UAB here instead of the platinum buffer since there would be limited plasma matrix remaining with the dilution. Samples and standards were mixed with the magnetic beads on a 96-well plate for 2 or 24 hours at ambient temperature on a shaker at 600 RPM (800 RPM for the first minute); this shaking condition applies to all subsequent incubation steps. Initially, we used a 2 h incubation period at this step with 7 standards, which is the recommended protocol. However, in later experiments, we extended the incubation time to 24 h at 5°C with an additional serial dilution to have 8 standards to improve protein capture with low concentration analytes of interest, per the recommendation from the vendor, and improve user workflow by splitting the assay into two days. Detection antibody mix was then added and incubated for 30 min, followed by addition of streptavidin-PE and a final 30 min incubation. Plates were washed twice with wash buffer between each step. After a final wash, 1X wash buffer was added to each well and the plate was shaken for 5 min before reading on the Luminex 200 with a minimum of 50 beads per analyte per sample, a 60 s timeout, and a DD gate of 7,500-25,000.

The raw data were analyzed using the ProcartaPlex Analysis App (Thermo Fisher Scientific) using a five-parameter logistic standard curve, and the concentration and/or fold change were plotted in Prism 10 (GraphPad). Conditions where the bead count was below 30 beads were excluded. When the reported concentration was listed as either <OOR (outside of range) or the value was below the lower limit of quantitation (LLOQ), the value was replaced with the LLOQ.

**Impact of pre-loaded protein stabilizer on blood hemolysis levels at increased time and temperature in capillary blood collected via Tasso mini device (pilot study shown in Figure 4b).** 1 healthy participant (female, P5) was recruited for this study. The participant collected capillary blood using six Tasso mini devices with EDTA-coated BD microtainers, where three BD microtainers were pre-loaded with 50 µL of Protein Plus and three BD microtainers had no stabilizer added. The collected blood was pooled amongst similar conditions before splitting them into 100 µL aliquots in 0.6 mL centrifuge tubes and incubating them in duplicate at various times (72 h, 120 h) and temperatures (25°C, 35°C), all with a 0 h control. Following the incubation, the plasma samples were processed and photographed as described above before storing the samples at -80°C.

**Method optimization to reduce sample matrix effects on immunoassay readout.** 11 healthy participants (8 female and 3 male, P1-8 and P13-15) were recruited for this study. Each participant collected capillary blood using three Tasso mini devices with EDTA BD microtainers pre-loaded with 50 µL of Protein Plus before pooling the blood from all three Tasso devices, collecting the plasma as described in the main manuscript, and stored at -80 C. We analyzed the plasma samples from one participant (noted as "P1") and pooled the plasma samples from the 10 remaining participants (noted as "Pooled") to account for variation between participants. To test the impact of the complex plasma matrix, we performed spike and recovery experiments for both P1 and Pooled samples by spiking cytokine standards at varied concentrations in different diluents: platinum buffer only, 1:4 plasma diluted in platinum buffer, and 1:8 plasma diluted in platinum buffer. The assay standards were prepared using platinum buffer and serially diluted using a 1:4 dilution to make 7 standards total, where standard 1 is the most concentrated. Standards 1-5 were spiked into the plasma samples (with and without dilution in platinum buffer) at a 1:10 dilution to create a standard curve (Table S6). Protein concentrations were measured by combining four ProcartaPlex simplex kits (Thermo Fisher Scientific) for IL-6, TNFa, CCL2, and CD40L according to the protocol described in the manuscript (Figure S1, CD40L data not shown). Each sample was measured in duplicate.

### III. SUPPLEMENTAL FIGURES

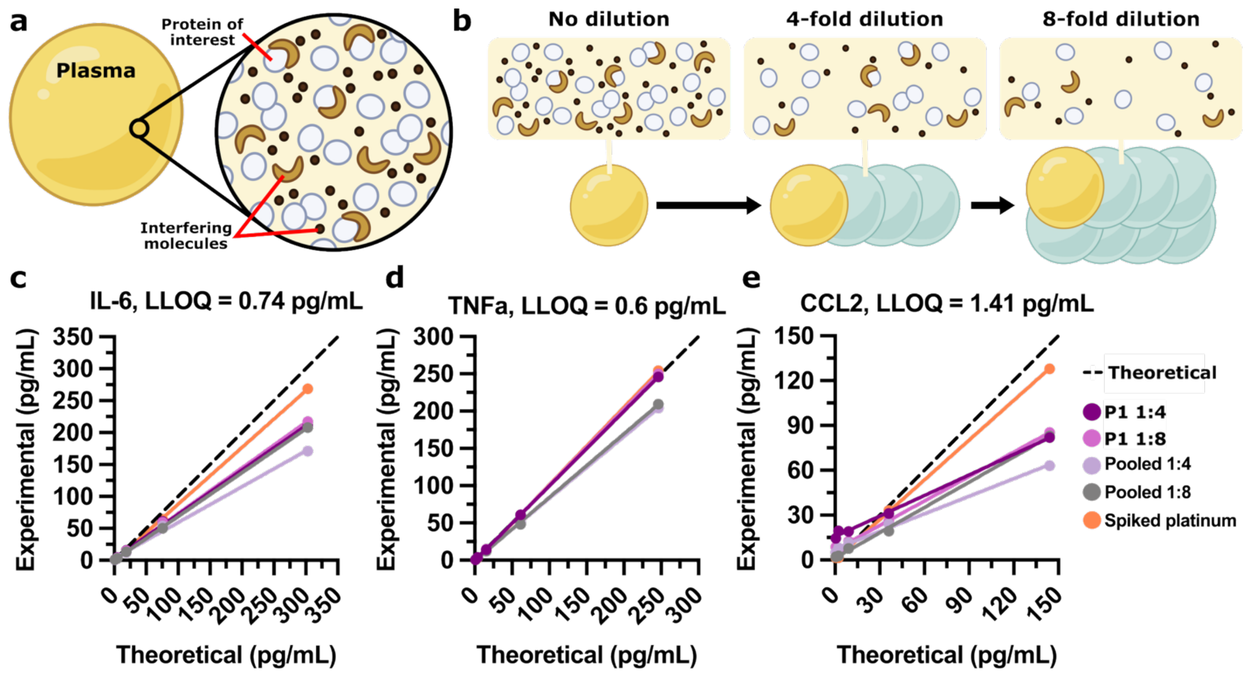


**Figure S1. Spike and recovery of proteins in complex plasma matrix improved by plasma dilution.** (a) Plasma is the liquid portion of anticoagulated blood and contains proteins of interest and other molecules that may interfere with downstream protein quantification (b) Interference from the complex plasma matrix can be reduced by diluting the sample. (c-e) Plasma from a single healthy participant (P1) and pooled from 10 healthy participants (Pooled, P2-8 & P13-15) collected via Tasso device was diluted in platinum buffer at either a 1:4 or 1:8 ratio. Assay standards were spiked into the plasma at various concentrations, with spiked platinum buffer as a control. The experimentally measured protein concentration was compared to the predicted (theoretical) concentration for (c) IL-6, (d) TNFa, and (e) CCL2. Each dot represents an average between two samples. The lower limit of quantitation (LLOQ) is noted above each graph.

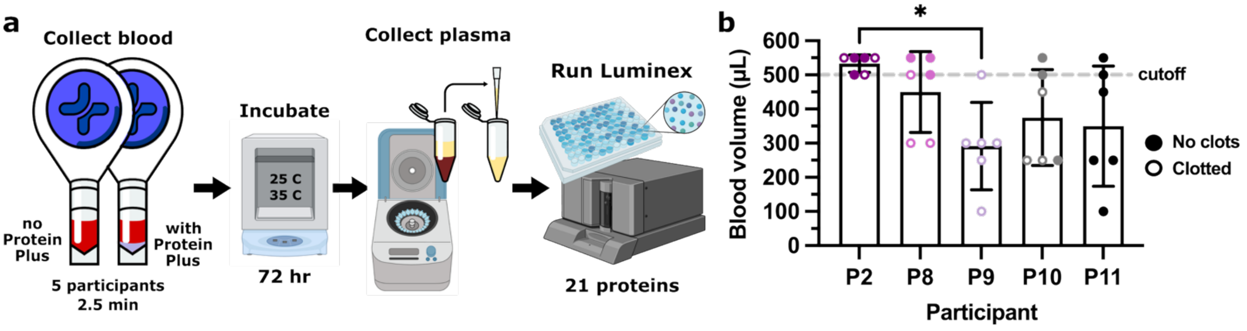


**Figure S2. Tasso temperature stress experimental workflow and blood volume.** This figure provides additional information for the experiment shown in Figure 5. (a) Capillary blood was collected via Tasso mini device into EDTA-coated BD microtainers either with no stabilizer (n = 3 Tassos pooled) or with 20 µL of Protein Plus (PP) stabilizer pre-loaded (n = 3 Tassos pooled) from 5 participants (P2, P8-11). After collection, the blood was incubated for 72 h at 25 C and 35 C before the plasma was isolated and stored at -80 C for at least two days before protein quantification via Luminex assay. (b) Estimated whole blood volume for all 5 donors. The volume cutoff (500 µL) is noted with a grey dashed line. Each point represents blood collected from a single Tasso. Results were compared using a one-way ANOVA with Tukey post-hoc tests ($n = 5$). All comparisons are not significant unless noted otherwise. * indicates $p < 0.05$.

### IV. SUPPLEMENTAL TABLES

**Table S1. Participant numbers and 3-letter codes for raw data interpretation.** Each participant number and code corresponds to a single participant and was kept constant throughout the manuscript.

| Participant number | Participant code |
|---|---|
| P1 | BDS |
| P2 | DKM |
| P3 | BRE |
| P4 | RJT |
| P5 | MYB |
| P6 | CWX |
| P7 | ZYT |
| P8 | GQS |
| P9 | PXN |
| P10 | TRJ |
| P11 | OPQ |
| P12 | UJB |
| P13 | PRX |
| P14 | DQX |
| P15 | BDZ |

**Table S2. Capillary blood collection time and clotting**

| Participant number | Tasso device (+ / mini) | Condition | Collection time (s) | Time to first blood (s) | Blood volume (µL) | Clots observed (Y / N) |
|---|---|---|---|---|---|---|
| P1 | + | 3-5 min | 198 | 28 | 1070 | Y |
| | + | 2.5 min | 152 | 31 | 600 | N |
| | + | 1.5 min | 91 | 34 | 330 | N |
| | + | Clotted control | 302 | 37 | N/A* | Y |
| | mini | 3-5 min | 302 | 33 | 940 | Y |
| | mini | 2.5 min | 152 | 33 | 580 | Y |
| | mini | 1.5 min | 93 | 48 | 230 | N |
| P2 | + | 3-5 min | 184 | 31 | 1000 | Y |
| | + | 2.5 min | 150 | 26 | 1050 | N |
| | + | 1.5 min | 92 | 31 | 570 | N |
| | + | Clotted control | 179 | 38 | N/A* | Y |
| | mini | 3-5 min | 301 | 60 | 730 | Y |
| | mini | 2.5 min | 152 | 19 | 1030 | Y |
| | mini | 1.5 min | 91 | 22 | 470 | N |

| | | | | | | |
|---|---|---|---|---|---|---|
| P3 | + | 3-5 min | 311 | 57 | 750 | Y |
| | + | 2.5 min | 147 | 33 | 810 | N |
| | + | 1.5 min | 88 | 24 | 550 | N |
| | + | Clotted control | 301 | 73 | N/A* | Y |
| | mini | 3-5 min | 301 | 44 | 570 | Y |
| | mini | 2.5 min | 151 | 48 | 380 | N |
| | mini | 1.5 min | 90 | 35 | 280 | N |
| P4 | + | 3-5 min | 180 | 24 | 1020 | N |
| | + | 2.5 min | 154 | 51 | 320 | N |
| | + | 1.5 min | 93 | 25 | 440 | N |
| | mini | 3-5 min | 304 | 52 | 640 | Y |
| | mini | 2.5 min | 152 | 28 | 1050 | N |
| | mini | 1.5 min | 92 | 29 | 425 | N |
| P5 | + | 3-5 min | 304 | 116 | 360 | Y |
| | + | 2.5 min | 152 | 56 | 420 | N |
| | + | 1.5 min | 93 | 72 | 225 | N |
| | mini | 3-5 min | 302 | 56 | 520 | Y |
| | mini | 2.5 min | 153 | 47 | 445 | N |
| | mini | 1.5 min | 93 | 70 | 175 | N |
| P6 | + | 3-5 min | 241 | 54 | 1120 | N |
| | + | 2.5 min | 160 | 22 | 1100 | N |
| | + | 1.5 min | 100 | 30 | 440 | N |
| | mini | 3-5 min | 306 | 36 | 600 | Y |
| | mini | 2.5 min | 158 | 133 | 250 | N |
| | mini | 1.5 min | 105 | 70 | 405 | N |
| P7 | + | 3-5 min | 234 | 32 | 1145 | Y |
| | + | 2.5 min | 135 | 23 | 1100 | N |
| | + | 1.5 min | 93 | 17 | 525 | N |
| | mini | 3-5 min | 302 | 144 | 650 | Y |
| | mini | 2.5 min | 152 | 62 | 305 | N |
| | mini | 1.5 min | 92 | 29 | 315 | N |

*N/A = not applicable. Volume not measurable via pipette when fully clotted

### Table S3. Blood and stabilizer volumes for varied ratios

| Blood volume (µL) | Protein Plus volume (µL) | Ratio |
|---|---|---|
| 300 | 0 | N/A |
| 300 | 10 | 1:30 |
| 300 | 15 | 1:20* |
| 300 | 20 | 1:15 |
| 300 | 30 | 1:10 |
| 300 | 60 | 1:5 |
| 300 | 100 | 1:3 |

*Recommended ratio

### Table S4. Capillary blood with and without stabilizer under shipping times and temperatures

| Participant number | Condition | Collection time (s) | Blood volume (µL) | Clots observed (Y / N) |
|---|---|---|---|---|
| P2 | no PP_1 | 132 | 550 | Y |
| | no PP_2 | 66 | 550 | N |
| | no PP_3 | 102 | 550 | Y |
| | with PP_1 | 92 | 500 | N |
| | with PP_2 | 90 | 550 | N |
| | with PP_3 | 147 | 500 | Y |
| P8 | no PP_1 | 150 | 500 | Y |
| | no PP_2 | 93 | 550 | N |
| | no PP_3 | 139 | 550 | Y |
| | with PP_1 | 103 | 500 | N |
| | with PP_2 | 150 | 300 | Y |
| | with PP_3 | 150 | 300 | Y |
| P9 | no PP_1 | 150 | 250 | Y |
| | no PP_2 | 150 | 500 | Y |
| | no PP_3 | 150 | 100 | Y |
| | with PP_1 | 150 | 300 | Y |
| | with PP_2 | 150 | 300 | Y |
| | with PP_3 | 150 | 300 | Y |
| P10 | no PP_1 | 150 | 500 | N |
| | no PP_2 | 150 | 450 | Y |
| | no PP_3 | 150 | 250 | Y |
| | with PP_1 | 150 | 250 | Y |
| | with PP_2 | 150 | 550 | N |
| | with PP_3 | 150 | 250 | N |
| P11 | no PP_1 | 150 | 250 | N |
| | no PP_2 | 150 | 500 | N |
| | no PP_3 | 150 | 450 | N |
| | with PP_1 | 160 | 100 | N |
| | with PP_2 | 150 | 250 | N |
| | with PP_3 | 90 | 550 | N |

### Table S5. Catalogue and lot numbers for Luminex kits

| Experiment | Kit | Catalogue number | Kit lot number | Standard lot number* |
|---|---|---|---|---|
| Blood collection time and clotting (Figure 2) | 19plex | PPX-19-MXZTGNT | 450767-000 | 389247-000<br>415041-000<br>389026-000<br>397396-000<br>420280-000<br>422159-000 |
| | Coagulation 3 | EPX04A-10825-901 | 450761-003 | 451462-000 |

| | | | | |
|---|---|---|---|---|
| Blood with and without stabilizer at shipping times and temperatures (Figure 5) | 20plex | PPX-20-MX9HMRW | 463248-000 | 415001-000<br>389247-000<br>319310-000<br>389026-000<br>415039-000<br>397396-000<br>420280-000<br>422159-000 |
| | CRP simplex | EPX01A-10288-901 | 450887-002 | 415041-000 |
| Plasma sample matrix effects (Figure S1) | IL-6 simplex | EPXS010-10213-901 | 426452-006 | 389820-000 |
| | TNFa simplex | EPXS010-10223-901 | 426489-005 | 389820-000 |
| | CCL2 simplex | EPXS010-10281-901 | 431783-006 | 436922-000 |
| | CD40L simplex | EPX01A-10239-901 | 444924-002 | 397396-000 |

*****Corresponds to each standard mix tube provided by each Luminex kit

### Table S6. Concentrations of spiked-in proteins for matrix effects experiment (Figure S1)

| Protein | Standard number | Initial concentration (pg/mL)* | Estimated concentration in sample (pg/mL)** |
|---|---|---|---|
| IL-6 | S1 | 3030.00 | 303.00 |
| | S2 | 757.50 | 75.75 |
| | S3 | 189.38 | 18.94 |
| | S4 | 47.34 | 4.73 |
| | S5 | 11.84 | 1.18 |
| TNFa | S1 | 2460.00 | 246.00 |
| | S2 | 615.00 | 61.50 |
| | S3 | 153.75 | 15.38 |
| | S4 | 38.44 | 3.84 |
| | S5 | 9.61 | 0.96 |
| CCL2 | S1 | 1440.00 | 144.00 |
| | S2 | 360.00 | 36.00 |
| | S3 | 90.00 | 9.00 |
| | S4 | 22.50 | 2.25 |
| | S5 | 5.63 | 0.56 |
| CD40L | S1 | 8050.00 | 805.00 |
| | S2 | 2012.50 | 201.25 |
| | S3 | 503.13 | 50.31 |
| | S4 | 125.78 | 12.58 |
| | S5 | 31.45 | 3.14 |

*Standard 1 was serially diluted at a 1:4 ratio to make standards 2-7.

**Standards 1-5 were spiked into the buffer and plasma samples at a 1:10 ratio.
